# Flavodiiron protein activity outcompetes cyclic electron transport when expressed in angiosperm *Nicotiana tabacum*

**DOI:** 10.1101/2025.01.23.634494

**Authors:** Nicholas Rizzetto, Eleonora Traverso, Andrea Sabia, Filippo Fiorin, Carlotta Francese, Livio Trainotti, Tomas Morosinotto, Alessandro Alboresi

**Affiliations:** Department of Biology, University of Padova, Italy; DAFNAE, University of Padova, Italy

## Abstract

In conditions of excess illumination, alternative electron transport pathways in the thylakoid membranes protect the photosynthetic apparatus against damage from eventual over-reduction. Two main pathways downstream of photosystem I (PSI) enable alternative electron flow, mitigating PSI acceptor-side limitation, while contributing to ATP biosynthesis without reducing NADP^+^ to NADPH: cyclic electron transport (CET) and pseudo-cyclic electron transport (PCET). Flavodiiron proteins (FLV) are crucial enzymes in PCET, found in all photosynthetic organisms but lost during the evolution of angiosperms. The absence of FLV coding sequences in angiosperm genomes raises intriguing questions about their role and function in photosynthetic organisms. Previous studies utilizing heterologous expression have already demonstrated that FLV can function in angiosperms. In this study, *Physcomitrium patens* FLVA and FLVB coding sequences were stably expressed in wild-type *Nicotiana tabacum*, a model crop species. Transgenic lines exhibited significantly increased PCET rates, with FLV-dependent electron transport competing for electrons with CET, particularly under sudden increases in light intensity that limited acceptor side limitation. These findings indicate that FLVs are not only active but also play a critical role in protecting from over-reduction the photosynthetic apparatus of *Nicotiana tabacum* under fluctuating light conditions.

## Introduction

Photosynthetic organisms utilize sunlight to drive electron transfer from water to NADP^+^, producing NADPH, through the activity of two photosystems (PSII and PSI). This process also generates a transmembrane electrochemical gradient, leading to ATP synthesis. Together, NADPH and ATP sustains carbon dioxide (CO_2_) fixation and several other metabolic reactions (Stirbet et al. 2020).

In natural environments plants live in dynamic conditions, including dark-to-light transitions, variable light intensities, and heterogeneous water and nutrient supply. These factors can disrupt the balance between excitation energy availability and the utilization of photosynthesis end-products, potentially leading to over-reduction of the electron transport chain. In the worst-case scenario, excess electrons may react with oxygen generating reactive oxygen species (ROS) that can cause irreversible damage to the photosynthetic apparatus (Joliot and Johnson 2011). Mechanisms regulating photosynthesis are crucial to respond to such imbalances, with a major impact on photosynthetic efficiency. Optimizing these mechanisms can also represent a potential strategy to enhance biomass accumulation under field conditions (Kromdijk et al. 2016; De Souza et al. 2022).

Photosynthetic organisms have evolved various mechanisms to dissipate excess energy, including non-photochemical-quenching (NPQ), cyclic electron flow (CEF) and pseudo-cyclic electron flow (PCEF). CEF and PCEF operate downstream of PSI, decoupling ATP synthesis from the one of NADPH production and protecting PSI from over-reduction (Alboresi et al. 2019b; Ruban and Wilson 2021; Leister 2023).

The two known CEF mechanisms recycle electrons from ferredoxin (Fd) back to plastoquinone (PQ) pool. The first is mediated by the chloroplast NADPH dehydrogenase-like (NDH) complex, homologous of the mitochondrial Complex I (Joliot and Johnson 2011). The second involves the PGR5/PGRL1 system, which facilitates electron transport from PSI back to the PQ pool (Munekage et al. 2002; DalCorso et al. 2008). Although PGR5 and PGRL1 proteins are clearly involved in regulating CEF (Munekage et al. 2002; Johnson et al. 2014), the precise mechanism by which the electrons are transferred from PSI acceptors to the PQ pool remains under debate (Nawrocki et al. 2019).

PCEF keeps PSI in an oxidized state by transferring P700-derived electrons to molecular oxygen, converting it into H_2_O (Messant et al. 2021). Since the electrons originate from H_2_O by the activity of PSII oxygen-evolving complex and ultimately reduce oxygen to water through PCEF, this process is also often termed the “water-water cycle”. PCEF contributes to ΔpH generation but does not support NADPH synthesis and is mediated by two distinct mechanisms: the Mehler reaction and flavodiiron proteins (FLVs).

Mehler reaction detoxifies the superoxide anion produced in PSI when molecular oxygen reacts with electrons from Fd or Fe-S clusters (Foyer and Hanke 2022). Superoxide is first converted into hydrogen peroxide by superoxide dismutase and subsequently scavenged into water molecules by ascorbate-peroxidase (Asada 2000).

In the stroma, FLVs accept electrons downstream of PSI to reduce molecular oxygen into H_2_O. These proteins belong to the multigenic flavodiiron protein (FDP) family, found across a diverse range of organisms (Folgosa et al. 2018). In photosynthetic organisms, FLVs consist of three functional domains: an N-terminal metallo-β-lactamases with a diiron-binding center followed by a domain able to interact with a flavin mononucleotide (FMN) and a final C-terminal domain with NADPH:flavin oxidoreductase activity (Folgosa et al. 2018; Alboresi et al. 2019a; Beraldo et al. 2024).

FLVs function as safety valves for PSI over-reduction in cyanobacteria (Allahverdiyeva et al. 2013), algae (Chaux et al. 2017; Shimakawa et al. 2022), non-vascular plants (Gerotto et al. 2016; Shimakawa et al. 2017) and gymnosperms (Ilík et al. 2017; Bag et al. 2023). They play a critical role during dark-to-light transitions and under fluctuating light intensities, when the Calvin-Benson cycle cannot fully utilize the reducing equivalents produced by the linear electron transport chain (Storti et al. 2020b). Their activity is negligible during steady-state photosynthesis.

Mutant lines deficient in FLV accumulation exhibit pronounced light sensitivity, highlighting the critical role of FLVs in photoprotection across various organisms. However, FLVs were lost during angiosperm evolution (Hanawa et al. 2017; Ilík et al. 2017) and are absent in certain eukaryotic algae, such as diatoms (Shimakawa et al. 2019).

One of the reasons why angiosperms have not conserved FLVs may be attributed to the increased efficiency of other electron sinks such as cyclic electron pathway (Yamamoto et al. 2016), the Calvin-Benson cycle and photorespiration (Hanawa et al. 2017; Shikanai and Yamamoto 2017).

Attempt to express heterologous cyanobacterial or moss *Physcomitrium patens* FLV genes in angiosperms showed that these proteins can function as electron acceptors downstream of PSI (Yamamoto et al. 2016; Gómez et al. 2018; Wada et al. 2018). Their expression has been shown to partially rescue the phenotype of cyclic electron flow (CEF) knock-out mutants (Yamamoto et al. 2016; Wada et al. 2018). In tobacco lines expressing *Flv1-Flv3* Synechocystis genes, photosynthetic performance under steady-state illumination remained similar to wild-type (WT) plants, but these lines exhibited a faster recovery of photosynthetic parameters during dark-to-light transitions (Gómez et al. 2018). Similarly, *Physcomitrium patens FLVA-FLVB* expression in WT *Arabidopsis thaliana* and *Oryza sativa* significantly increased PCET rates under fluctuating light conditions but had no measurable impact during steady-state photosynthesis (Yamamoto et al. 2016; Wada et al. 2018).

These initial studies, while demonstrating that FLVs can function in angiosperms and contribute to photoprotection under specific conditions, left several important questions unresolved. One critical uncertainty is whether FLV activity *in vivo* offers a measurable advantage compared to WT plants, particularly in terms of photosynthetic performance or stress tolerance. Additionally, it remains unclear under which environmental or physiological conditions FLVs might enhance tolerance to excess light, such as fluctuating light intensities, prolonged high-light exposure, or rapid dark-to-light transitions. To further investigate the impact of FLV in angiosperms, we expressed *FLVA-FLVB* from *P. patens* in tobacco. Our results confirmed that FLVs are active in this system and capable of supporting high rates of electron transport, with kinetics comparable to those observed in *P. patens*. FLV activity was found to compete with CET for electrons, supporting the hypothesis that CET compensates for the absence of FLVs in angiosperms. Additionally, PCET is also shown to be effective in mitigating PSI acceptor-side limitation upon exposure to light fluctuations, demonstrating a significant role in enhancing photoprotection.

## Materials and methods

### Plant material and growth conditions

*Nicotiana tabacum* cv. Samsung NN, WT and FLV-expressing plants were sowed on watered filter paper and after germination plantlets were transferred in germinators for one week and finally they were planted into single pots for all the analyses. Plants were grown in a growth chamber at 25°C, 45-65% air humidity under a photoperiod of 16-h day (100 μmol m^-2^ s^-1^) and 8-h night. To test photosynthetic efficiency under fluctuating light conditions, 3 weeks-old plants were positioned under LED lamp (Photon System Instruments; SL 3500) and subjected to 4.5 minutes at 50 μmol m^-2^ s^-1^ followed by 30 seconds at 1000 μmol m^-2^ s^-1^. The photoperiod was 16-h of fluctuating light and 8-h night. To increase the stress related to light fluctuation, this treatment was performed in a growth chamber at 16°C.

*Physcomitrium patens* ssp. Gransden WT and *flv* KO (Gerotto et al. 2016). All plants were routinely propagated as previously described on PpNH_4_ rich medium at 24°C under long-day conditions (16-h day/8-h night). For photosynthetic measurements and for plant growth quantification the individual lines were cultivated photoautotrophically on PpNO_3_ minimal medium (Ashton et al. 1979; Storti et al. 2019).

### Molecular cloning and plant transformation

The introduction of the two *P. patens FLVA* and *FLVB* genes in *N. tabacum* was achieved by expressing the *FLVA-2A-FLVB* sequence (Yamamoto et al. 2016) under the control of the 35S constitutive promoter. *FLVA-2A-FLVB* gene (4182 bp) was subcloned in pDONR221 and then transferred into the pBinAR expression vector (Hofgen and Willmitzer 1988) using the Gateway cloning system to obtain pBinAr-35S::FLVA-2A-FLVB vector. *Agrobacterium tumefaciens* strain LBA4404 was used for the stable transformation of *N. tabacum* plants (Fisher and Guiltinan 1995). Young, healthy, and undamaged leaves were collected from *in vitro*-grown WT tobacco plants and soaked in a suspension of *A. tumefaciens* cells (OD_600_ ∼0.8) in MS salts without vitamins (4.3 g/L), sucrose (30 g/L), 6-BAP (1 mg/L) and Gamborg B5 medium (1 ml/L of 1000x stock solution) for 10 minutes to allow the infection cycle. Leaves were then placed on TAB1 co-cultivation medium (MS salts with vitamins 4.4 g/L, sucrose 30 g/L, 6-BAP 1 mg/L, IAA 0.2 mg/L, plant agar 5 g/L, pH 5.7) and incubated in the dark at 23°C for 48 h. Leaves were then transferred to plates containing TAB2 medium (TAB1 medium supplemented with kanamycin 200 mg/L and cefotaxime 500 mg/L) and incubated at 23°C with a 16-hour light/8-hour dark photoperiod. The TAB2 medium was refreshed every 15 days until callus formation occurred. Once shoots of 1-3 cm in length emerged from the callus masses, independent clones were cut off and placed in Magenta boxes containing TAB3 medium (MS salts with vitamins 4.4 g/L, sucrose 30 g/L, plant agar 6 g/L, kanamycin 200 mg/L and cefotaxime 500 mg/L, pH 5.7) for rooting. After ∼20 days, plants were analyzed by PCR on genomic DNA to confirm the presence of the transgene and then transferred to soil. The selected T_0_ plants were self-crossed to obtain the corresponding T_1_ and T_2_ lines. Segregation analysis of the transgene was conducted by sowing pre-sterilized T_2_ seeds in MS 1/2 medium supplemented with kanamycin 200 mg/L (Fig. S2). The plates were incubated in a growth chamber at 25°C (16-hour light/8-hour dark photoperiod) for 15 days, time after which germinated plants, resistant to kanamycin selection, showed green developed cotyledons.

### Gene expression analysis

RNA was purified from leaf disks with TRI Reagent® (Sigma-Aldrich) and used as template for cDNA synthesis to verify FLVA, FLVB and H2A expression. The primers used for PCR are the following FLVAf: 5’-GATGCCCCGGGATGGAGACGATGTTGGTG-3’, FLVAr: 5’-CACCCTAACATCTGCCTCCT-3’, FLVBf: 5’-CTGTCAAAGCCAAGCAACCC-3’, FLVBr: 5’-AGGACTTCCATTTGCCCCAG-3’, H2Af: 5’-GAGTTACCATCGCAAGCGGA-3’ and H2Ar: 5’-ATGCCTTCTCCTCAGCTGCA-3’.

### Immunoblot analysis

Total extracts were obtained by grinding leaf disks in 150 μL sample buffer (50 mM TRIS pH 6.8, 100 mM DTT, 2% SDS, and 10% glycerol) and centrifugated at 10,0000 g. An aliquot of the supernatant was used for chlorophyll extraction in acetone 80%. Chlorophyll content was quantified at the spectrophotometer (Cary Series UV-Vis, Agilent Technologies) applying the formula below (Porra et al. 1989):

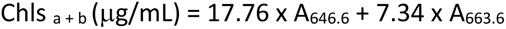

The supernatant was incubated at 100°C for 1 min before loading into SDS/PAGE. The quantity of samples to load was estimated in terms of equivalents of chlorophyll amount. For immunoblotting analysis, after SDS/PAGE, proteins were transferred to nitrocellulose membranes (Pall Corporation) and hybridized with specific commercial antibodies (PSAA, Agrisera, catalog numbers AS06 172) and homemade polyclonal antibodies (PsaD, γATPase, Cyt f, PsbS, D2, NDHH, RuBisCO and LHCII) and custom-made anti-FLVB. The proteins were detected with alkaline phosphatase-conjugated secondary antibody and colorimetric assay.

### Fluorescence and P700 Measurement with Dual-PAM

The redox changes of P700 were recorded using a Dual-PAM100 measuring system (Heinz Walz, Effeltrich, Germany) during 6 s long illumination (2000 μmol m^-2^ s^-1^). Before the measurement, moss samples and tobacco leaves were dark adapted for 30 min. Samples were measured under standard atmospheric conditions as well as anaerobic conditions. Anaerobiosis was induced by incubation of the samples in a plastic box filled with nitrogen enriched atmosphere for 15 to 20 min. The oxidation level of P700 was quantified following a repeated Saturating Pulse protocol (rSP) (Shimakawa et al. 2019). Leaves were dark-adapted for 30 minutes and then exposed to a first saturating pulse for the determination of the total oxidizable P700 (Pm) and then to 6 SPs (∼2000 μmol m^-2^ s^-1^) lasting 1 s applied every 10 s in presence of low actinic light (∼70 μmol m^-2^ s^-1^). The fraction P700^+^/total oxidizable P700 was calculated by integrating the P700^+^ during the last 1s-SP, normalized to the Pm. The effects of fluctuating light on photosynthesis were evaluated on 3-month-old dark-adapted plants. The light protocol consisted in 3 cycles of 5 min low actinic light (∼60 μmol m^-2^ s^-1^) followed by saturating actinic light for 1 min (∼1600 μmol m^-2^ s^-1^). The following PSI and PSII parameters were calculated: Y (I) as 1-Y (ND)-Y (NA), Y (NA) as (P_m_-P_m′_)/P_m_, Y (ND) as (P700_ox_/P_m_), Y (II) as (F_m′_-F)/F_m′_, 1-qL as qL = (F_m′_-F)/(F_m′_ F_0’_) × F_0’_/F, NPQ as (F_m_-_Fm′_)/F_m′_, where F_0_ and F_m_ represent the minimum and maximum chlorophyll fluorescence levels (Maxwell and Johnson 2000), whereas P_m_ and P_m′_ represent the P700 signals recorded before and after onset of a saturating pulse (Klughammer and Schreiber 1994). The electron transport rate around PSI and PSII was calculated as follows: ETRI as PAR x ETR-Factor x P_PS2_/P_PS1+2_ x Y (I) and ETRII as PAR x ETR-Factor x P_PS2_/P_PS1+2_ x Y (II), where ETR-Factor represents the sample absorptance and P_PS2_/P_PS1+2_ the distribution of absorbed PAR to PSII.

### Spectroscopic analyses with a Joliot-type spectrometer

Spectroscopic analyses were performed using a Joliot-Type spectrometer (JTS)-10 (Biologic). The change in absorption spectra of pigments due to electric potential gradient, known as electrochromic shift (ECS), was used to calculate the relative amount of PSI and PSII and the electron transport rates (ETR). For each sample ECS was measured after 30 min of dark adaptation at 520 nm and consecutively at 546 nm to subtract the field-independent contribution. Total number of functional PSI and PSII was determined by a xenon-induced single flash turnover and was then used to normalize ETR values. Plants were then exposed to 5 min of continuous light (940 μmol m^-2^ s^-1^) and during this time a dark-interval relaxation kinetic was applied: light was repeatedly switched off to calculate the non-photosynthetic contribution to ECS. ETR was calculated as the difference between the ECS slope in the light and the slope calculated in the dark (SD). The same measurements were carried out also in presence of 3-(3,4-dichlorophenyl)-1,1-dimethylurea (DCMU). DCMU was used to inhibit PSII and isolate the contribution of ETR around PSI to ECS. Before detection two 1 cm^2^ leaf samples were detached from each dark-adapted plant. One of them was incubated for 1 hour in paper soaked with DCMU 80 μm and the other one in deionized water as control.

## Results

### Isolation of Nicotiana tabacum lines expressing Physcomitrium patens FLVA and FLVB genes

The coding sequences of FLVA and FLVB from *Physcomitrium patens* were introduced into *Nicotiana tabacum* to investigate the role and functionality of these proteins in an angiosperm system. The open reading frame consisted of *FLVA* and *FLVB*, separated by the self-cleaving 2A-peptide derived from foot-and-mouth disease virus. This construct was designed to guarantee a stoichiometric production of FLVA and FLVB, that form an active heterocomplex, a strategy previously validated in *A. thaliana* and *O. sativa* (Yamamoto et al. 2016; Wada et al. 2018). The final FLVA-2A-FLVB sequence was cloned under the control of the constitutive 35S promoter and transformed into *N. tabacum* leaf disks, enabling the generation of transgenic tobacco lines for further characterization.

The functionality of FLVs can be assessed by measuring the redox kinetics of P700 in PSI under a short-pulse light illumination (20000 μmol m^−2^ s^−1^ for 6 seconds) where the protein impact can be evidenced by the ability to maintain PSI in a oxidized state (Ilík et al. 2017; Shimakawa and Miyake 2018). Given its reliability and efficiency, monitoring P700 oxidation kinetics was selected as a rapid screening method to identify functional FLVs in transgenic *Nicotiana tabacum* clones. The impact of FLV presence on these measurements is shown comparing *P. patens* WT and *flva/b* KO mosses. In both moss lines, the initial P700 oxidation results from PSI charge separation event followed by P700 reduction attributed to electrons extracted from water by PSII and reaching PSI (Ilík et al. 2017). During the light pulse, the WT P700 pool was oxidized again through the FLV-mediated electron transport downstream of PSI (Fig. 1). In contrast, P700 in *flva/b* KO mosses remained reduced throughout the light exposure (Fig. 1).

**Fig. 1.**
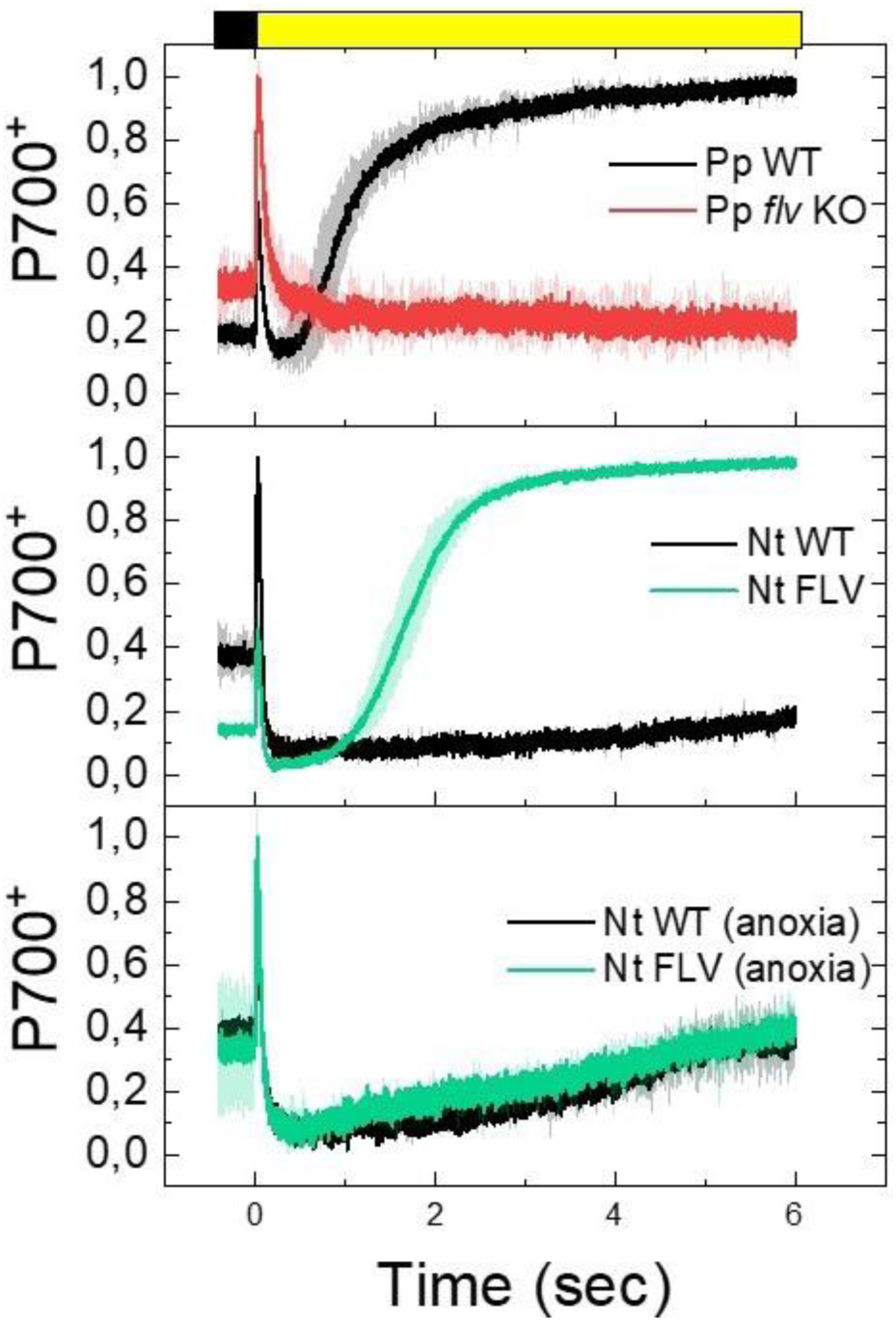
Redox kinetics of P700 upon dark-to-light transition in *Physcomitrium patens* protonemata cells and *Nicotiana tabacum* leaves. P700 redox kinetics were monitored *in vivo* during dark-to-light transitions in wild-type (WT) (black line) and *flva/b* KO lines (red line) of *P. patens* (upper panel), as well as in WT (black line) and representative transgenic *N. tabacum* leaves expressing moss FLVs (green line) (middle and lower panels). The kinetics of redox changes were measured *in vivo* upon exposure of dark-adapted sample (black bar) to actinic light (2000 mmol m^-2^ s^-1^; yellow bar). Light was switched on at time 0 and illumination lasted for 6 seconds. Samples were either measured under standard atmospheric conditions (top and middle panel) or anaerobic conditions (lower panel). Each curve is the mean of three independent biological replicates, with standard deviation shown as shaded areas.

In *N. tabacum*, WT plants exhibited a P700 oxidation profile comparable to that observed in *P. patens flva/b* KO lines, with P700 that remained reduced after the initial illumination (Fig. 1). On the contrary, some transformed *N. tabacum* lines exhibited a distinct P700 oxidation pattern resembling the one of WT mosses thus suggesting they expressed functional FLVs (Fig. 1). Since FLV activity is oxygen-dependent, an additional control experiment was performed using the same saturating light pulse protocol under anoxic conditions. In WT *N. tabacum* plants, the P700 oxidation profile remained unaltered regardless of oxygen availability (Fig. 1B-C). In contrast, the putative FLV-expressing lines lost their ability to keep P700 in an oxidized state under anoxia, exhibiting a WT-like behavior (Fig. 1B-C). This oxygen-dependent response confirmed the presence and activity of functional FLV proteins in these lines.

Further validation of FLV expression in *N. tabacum* transgenic clones was performed through immunoblot analysis, which detected the accumulation of *P. patens* FLVB in multiple lines. Among the transgenic *N. tabacum* lines showing positive results in both P700 oxidation kinetics and FLV protein accumulation, clones #1, #7 and #8 were selected for propagation to F2 generation and further investigations (Table S1).

RT-PCR showed that both FLVA and FLVB genes were expressed. (Fig. 2A) Immunoblot analysis confirmed that the accumulation levels of FLVB proteins were comparable among the selected lines (Fig. 2B). To assess whether the expression of FLV proteins impacted the photosynthetic machinery in transgenic *N. tabacum* lines, we analyzed the accumulation of key proteins associated with photosystems and electron transport. The accumulation of proteins constituting PSII (D2) and PSI (PSAA and PSAD) as well as major antenna proteins (LHCII) showed no evident alterations in FLV transgenic lines as compared to WT. The abundance of Cyt *f* and γ-ATPase also did not significantly change in FLV lines compared with WT (Fig. 2B). The levels of PsbS, the key activator of non-photochemical quenching (NPQ) of chlorophyll fluorescence at the level of PSII, also remained unchanged between WT and FLV lines (Fig. 2B). No significant changes in the accumulation of the RuBisCO enzyme or the NDH-H subunit were detected between WT and FLV lines (Fig. 2B). Overall, the accumulation of FLV proteins had thus no major effect on the composition of the *N. tabacum* photosynthetic apparatus. Since the three independent lines exhibited consistent behavior across all analyses, from here on the overall averaged data are presented for clarity.

**Fig. 2.**
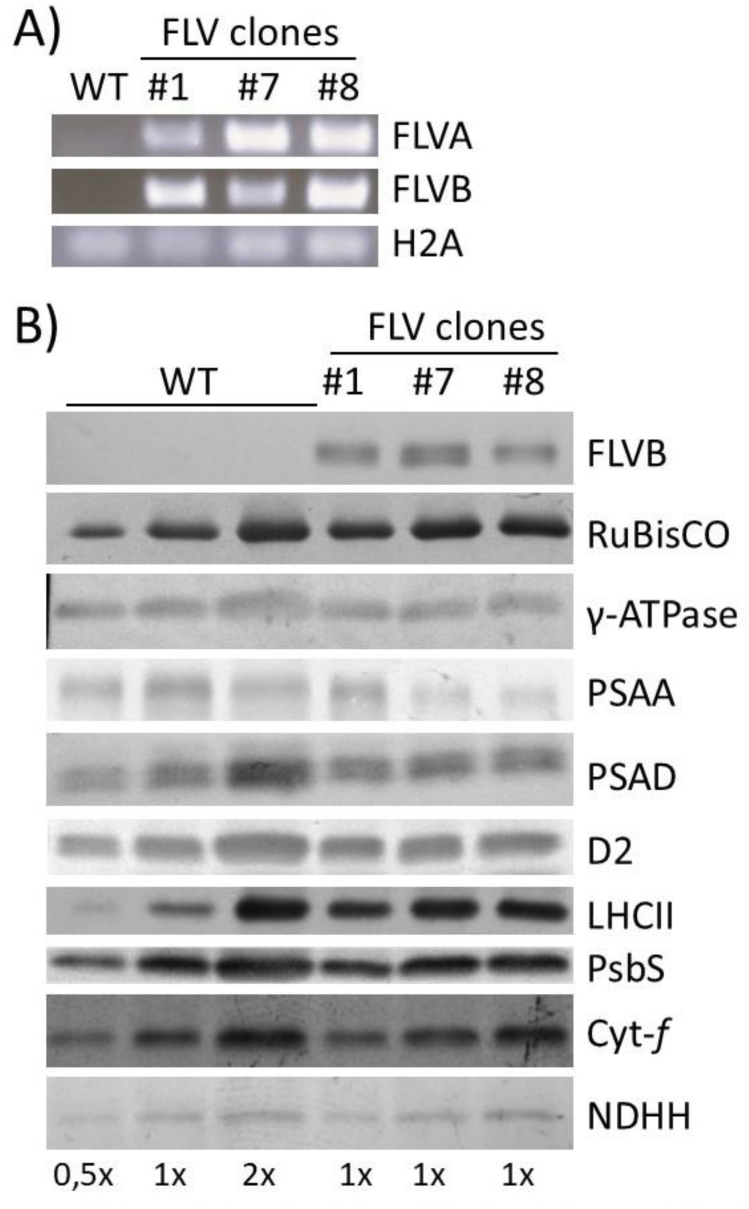
FLV expression and Immunoblot analysis of photosynthesis-related proteins in *N. tabacum* lines. A) The expression of FLVA and FLVB was verified via RT-PCR, H2A hystone served as the endogenous control. B) Immunoblot analysis was performed to detect photosynthesis related proteins in both wild-type (WT) and transgenic lines of *N. tabacum* expressing FLVA-2A-FLVB. The proteins analyzed included FLVB, RuBisCO, γ-ATPase, PSAA, PSAD, D2, LHCII, PsbS, Cytf, and NDHH. The detection was carried out using different chlorophyll equivalents as follows: 1× is equivalent to 2 µg of chlorophylls for the detection of FLVB, γ-ATPase, PSAA, PSAD, D2, PsbS, Cyt-f and NDHH. 1× is equivalent to 0.2 µg of chlorophylls for the detection of RuBisCO and LHCII. The amounts 0.5× and 2× represent half and twice the amount of 1×, respectively.

To validate FLV activity and confirm the initial screening results, we employed a quantitative protocol to determine the fraction of the fraction of oxidized P700 (P700^+^) relative to the total amount of photo-oxidizable P700. To this aim, plants were exposed to repetitive 1-second saturating pulses (rSPs) over a background of moderate light intensity between rSPs (Shimakawa and Miyake 2018) (Fig. 3A and Fig. S1). In WT plants PSI was oxidized at only 20% of its maximum capacity, while in FLV lines, 80% of PSI reaction centers were kept in an oxidized state during the last 1-second treatment (Fig. 3B and Fig. S1). This result confirmed that *P. patens* FLV expressed in *N. tabacum* was actively oxidizing P700.

**Fig. 3.**
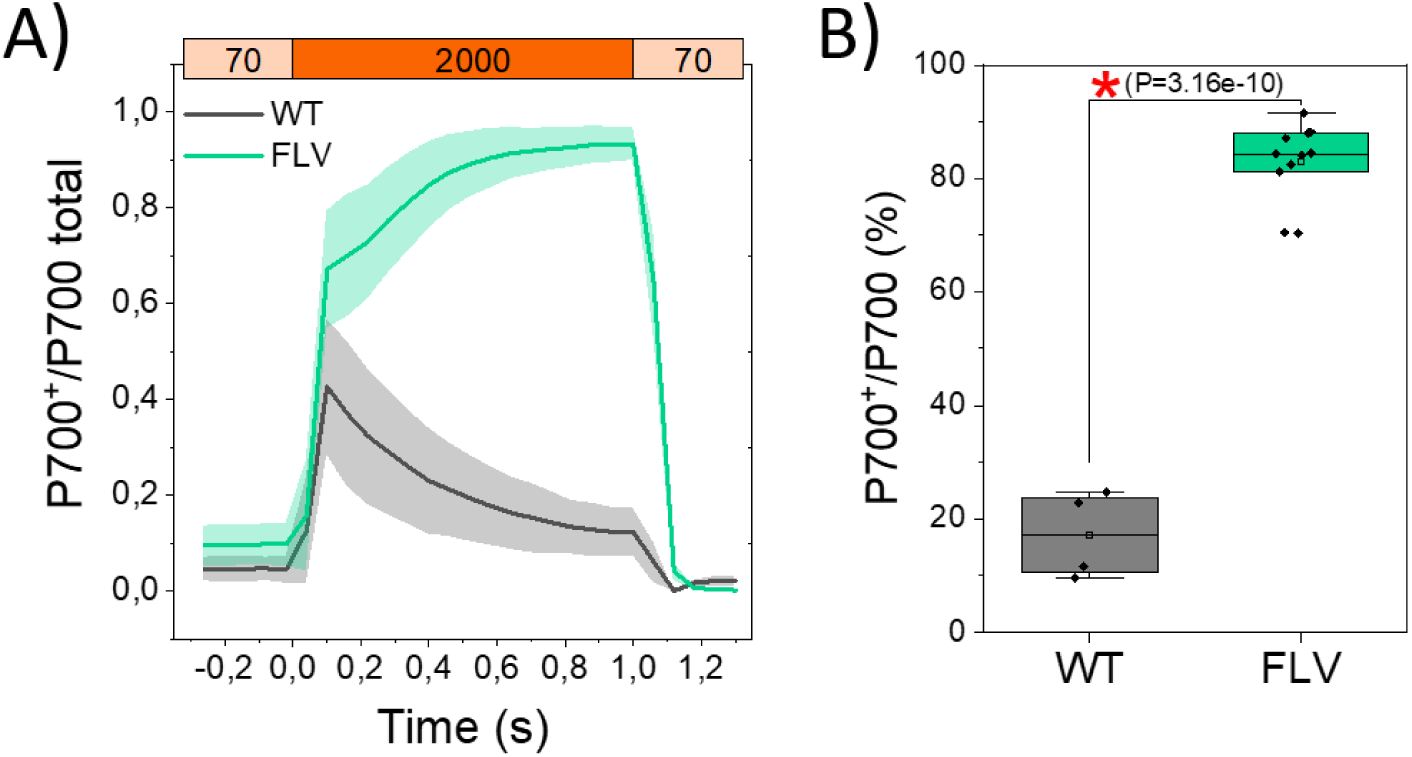
Oxidized P700 fraction in *Nicotiana tabacum* plants. A) Comparison between wild-type (WT) and the FLV-expressing lines, the kinetics of oxidized P700 (P700^+^) during illumination with a short-pulse light (SP: ∼2000 μmol m^−2^ s^−1^, 1 s). WT plants (black) and FLV-expressing lines (green) were subjected to SP in the presence of a background light of ∼70 μmol m^−2^ s^−1^. The relative P700^+^ amount is normalized to Pm, which represents the maximum oxidation level of P700. Standard deviation is indicated as a shadow. B) P700 oxidation index shown as percentage of P700^+^ over the total P700. The integration of the P700^+^ was calculated as the sum of relative P700^+^ values every 0.3 ms during SP illumination, which was then divided by the maximum integrated area. WT, n = 4 ± SD. FLV, n = 11 ± SD. A red asterisk indicates statistical significance (one-way ANOVA).

### FLV expression enhances the response of N. tabacum to fluctuating light conditions

In *P. patens* plants, FLV activity was indispensable in dark-to-light transitions and after a sudden increase in incident illumination (Gerotto et al. 2016; Storti et al. 2020b) and consistent results were found in all FLV expressing organisms (Allahverdiyeva et al. 2013; Chaux et al. 2017; Shimakawa et al. 2017). Therefore, we challenged *N. tabacum* plants with three cycles of limiting (low) and saturating (high) light intensities, monitoring different photosynthetic parameters (Fig.4 and Fig. S2).

**Fig. 4.**
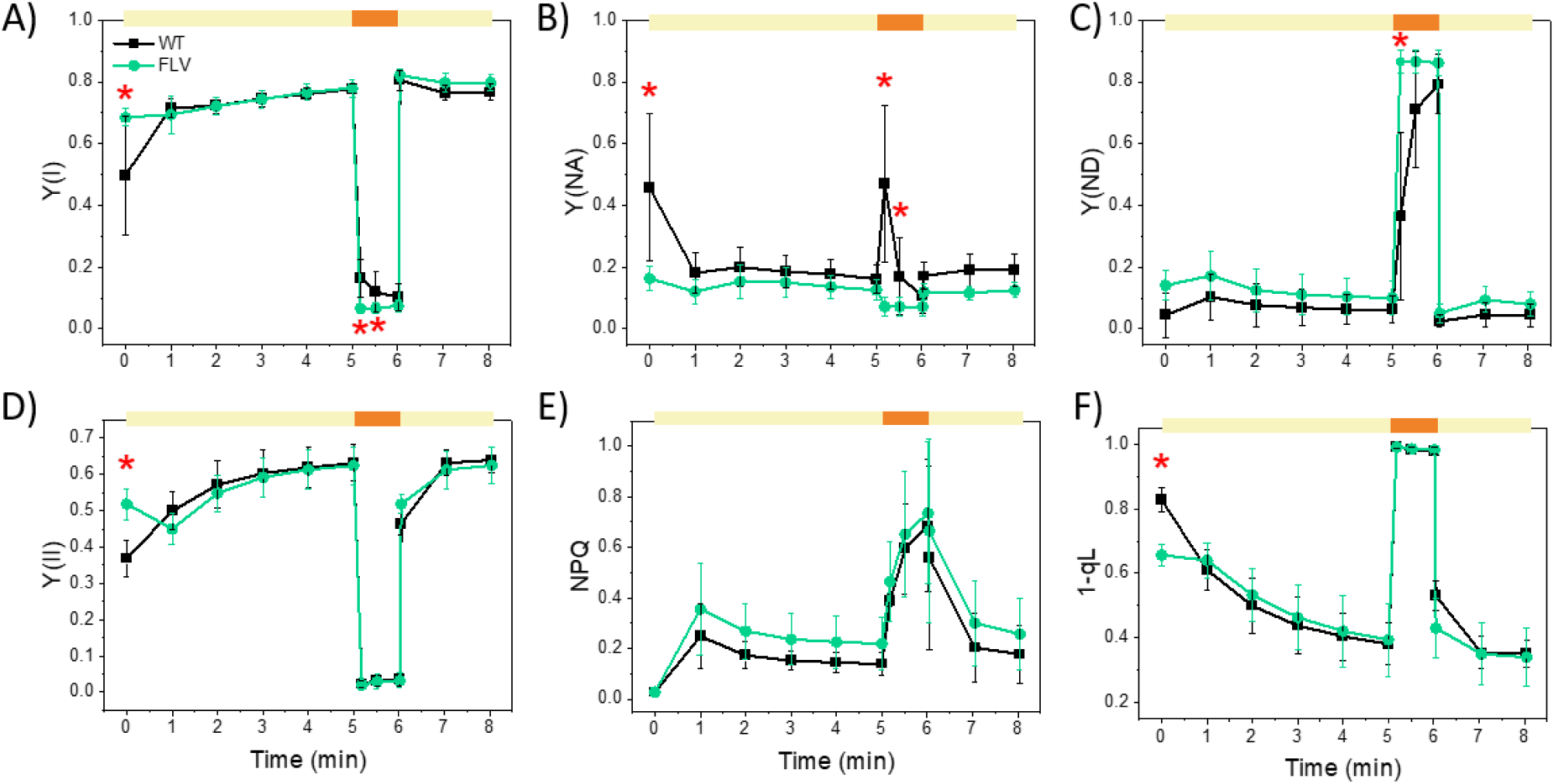
Effect of *P. patens* FLVs on photosynthetic parameters of *N. tabacum*. Photosynthetic efficiency was monitored during fluctuating light cycles for WT plants (black squares) and FLV-expressing lines (green circles): Y(I) (A), Y(NA) (B), Y(ND) (C), Y(II) (D), Y(NPQ) (E), and 1–qL (F). At time 0, after 40 min of dark adaptation, plants were treated with low actinic light (∼60 μmol m^−2^ s^−1^; yellow bars) for 5 min followed by saturating actinic light (∼1600 μmol m^−2^ s^−1^; orange bars) for 1 min. This cycle was repeated 2 more times as shown in Supplementary Figure S2. Data are averages ±SD, with statistical significance indicated by red asterisks (P < 0.01, one-way ANOVA). Data represents average values ±SD, n=7 for the WT, n=17 for FLV lines. Data are averages ±SD, with statistical significance indicated by red asterisks (P < 0.01, one-way ANOVA). WT plants: n = 7; FLV lines: n = 17.

All genotypes showed the same initial F_v_/F_m_ values (WT=0.78 ± 0.01; FLV lines=0.77 ± 0.01). During the initial transition from darkness to low light, WT plants showed a decrease of Y(I) and Y(II) that was recovered within a few minutes of illumination (Fig. 4A and D). This drop is likely the result of the transition from dark to light and indeed this effect was not present in the following low light cycles (Fig. S2), when plants were already light adapted. Interestingly, FLV-expressing lines showed higher YI and YII with respect to WT in this dark to low light transition (Fig. 4A and D). Upon exposure to a stronger light, after 5 minutes in LL, both Y(I) and Y(II) strongly decreased to ∼0.2, as result of the saturation of photosystem photochemical capacity (Fig. 4A and D). During the third cycle, Y(I) values during low-light periods were significantly higher in FLV lines (∼0.8) than in WT plants (∼0.73) (Fig. S2).

PSI activity and efficiency depends on acceptor or donor side limitation, estimated respectively as Y(NA) and Y(ND) parameters. WT plants displayed a steep and transitory Y(NA) increase at each transition from low-to-high light intensity (Fig. 4B and Fig. S2). By the end of the HL treatment, this limitation is recovered, and PSI is only limited from the donor side (Fig. 4C and Fig. S2). In FLV-expressing plants Y(NA) was instead constant and always very low (Fig. 4B and Fig. S2), suggesting that FLV were ready to receive electrons from PSI, working as additional acceptor. Because of the sustained flow of electrons towards FLVs during high light, PSI of transgenic lines was always limited at the donor side (Fig. 4C and Fig. S2).

Analyzing PSII specific parameters, no significant differences in NPQ (Fig. 4E and Fig. S2) and in the redox state of plastoquinone pool (1-qL; Fig. 4F and Fig. S2) were observed between WT and FLV-expressing lines, but at the very beginning of low light period for1-qL.

Estimation of photosynthetic electron transport rates (ETR) provided further insights into the functional differences between WT and FLV-transgenic lines, particularly during the transition from low to high light intensity (Fig.5 and Fig. S3). At the PSI level, the electron transport rate (ETRI) in WT plants was significantly higher than in FLV-transgenic lines within the first few seconds of the low-to-high light transition (Fig. 5A). At the PSII level, the estimated electron transport rate (ETRII) was comparable between WT and FLV-expressing plants across all light treatments (Fig. 5B). This consistency indicates that FLVs, despite their role as an alternative electron sink downstream of PSI, do not significantly affect the photochemical activity of PSII in these conditions. To assess cyclic electron flow (CEF) activity, we calculated the difference between ETRI and ETRII, as described previously (Huang et al. 2015). WT plants exhibited transiently higher CEF activity during the first 20 seconds of high light exposure compared to FLV-expressing lines (Fig. 5C and Fig. S3). This spike in CEF for WT suggested how this alternative electron flow played a major role in mitigating the over-reduction of PSI. Differently, FLV-expressing lines showed a reduced reliance on CEF during this transition (Fig. 5C and Fig. S3).

**Fig. 5.**
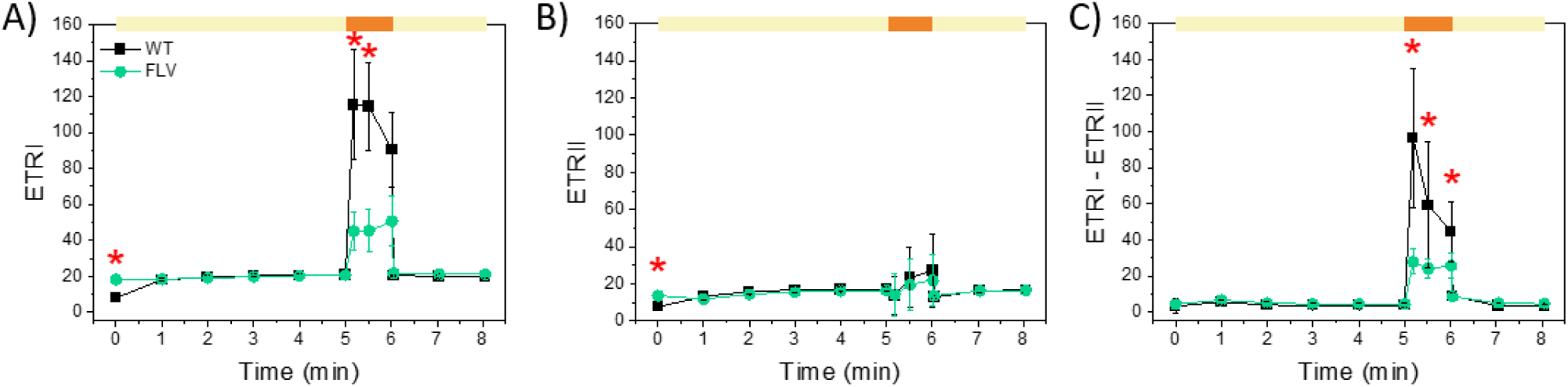
*P. patens* FLVs suppress cyclic electron transport in *N. tabacum*. Photosynthetic electron transport rates in WT (black squares) and FLV-expressing lines (green circles) were measured during fluctuating light cycles. ETRI (A), ETRII (B) and ETRI-ETRII (C) as a proxy for cyclic electron transport. At time 0, after 40 min of dark adaptation, plants were treated with low actinic light (60 μmol m^−2^ s^−1^; yellow bars) for 5 min followed by saturating actinic light (1600 μmol m^−2^ s^−1^; orange bars) for 1 min. This cycle was repeated 2 more times as shown in Supplementary Figure S3. Data are averages ±SD, with statistical significance indicated by red asterisks (P < 0.01, one-way ANOVA). WT plants: n = 7; FLV lines: n = 17.

### P. patens FLVs boost electron transport at light onset of in N. tabacum

To corroborate the estimated differences in ETRI and ETRII, photosynthetic electron transport was quantified by electrochromic shift (ECS) of carotenoid absorption through dark-interval relaxation kinetics (DIRK) analysis (Sacksteder and Kramer 2000). In dark-acclimated WT plants, ETR started at ∼20 electrons per second and remained stable during the first 20 seconds of illumination. Successively ETR steadily increased reaching a steady state after 4-5 minutes of illumination when carbon fixation was fully activated (Figure 6A and Fig. S4). In *N. tabacum* lines expressing FLV, the ETR was twice that of WT at the onset of actinic light (∼40 electrons per second; Fig. 6A and S4). This burst of activity was transient, persisting for only a few seconds. Subsequently, the ETR decreased to levels even lower than those of the wild type (WT). After approximately 5 minutes of light initiation, when steady-state photosynthesis was achieved, no statistically significant differences were detected among the genotypes. (Fig. 6A and Fig. S4).

**Fig. 6.**
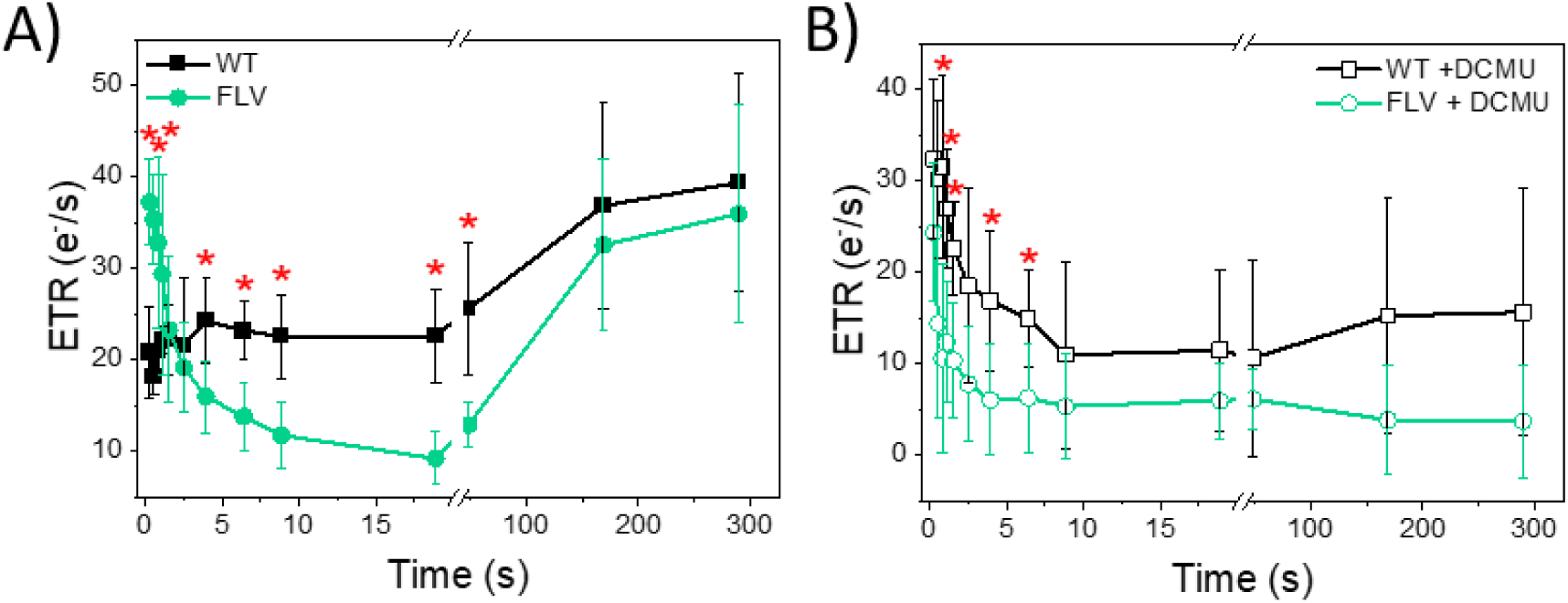
Photosynthetic electron transport in *N. tabacum* WT and FLV-expressing lines. (A) Total photosynthetic ETR measured in WT (black squares) and FLV-expressing lines (green circles) at 940 μmol m^−2^ s^−1^ actinic light, calculated from electrochromic shift signal. (B) Cyclic electron transport rate measured in the same samples treated with the PSII inhibitor 3-(3,4-dichlorophenyl)-1,1-dimethyl urea (DCMU). Data represents average values ±SD, n=6 for the WT, n=17 for FLV lines. Differences between WT and mutant plants were examined by one-way ANOVA; a red asterisk indicates statistical significance (P < 0.01).

To further dissect the impact of FLVs, DIRK analysis was conducted on leaf disks treated with the PSII inhibitor 3-(3,4-dichlorophenyl)-1,1-dimethylurea (DCMU), to specifically quantify CET around PSI (Fig. 6B and Fig. S4). WT plants showed high CET activity at the initiation of illumination followed by a gradual decline to a steady activity. Conversely, FLV-expressing lines consistently exhibited lower CET activity compared to WT throughout the first seconds of light treatment (Fig. 6B and Fig. S4).

### FLV expression increases tolerance to fluctuating light during growth

To elucidate the impact of FLVs on the photosynthetic activity during plant growth, WT and FLV expressing lines of *N. tabacum* were grown under constant control light intensity (CL) and the maximum efficiency of PSII and PSI was monitored as Fv/Fm and Pm (the maximal change in the P700 signal), respectively (Fig. 7 and Fig. S5). WT and FLV-expressing lines showed similar values both at the level of Pm (Fig. 7A) and Fv/Fm (Fig. 7B).

**Fig. 7.**
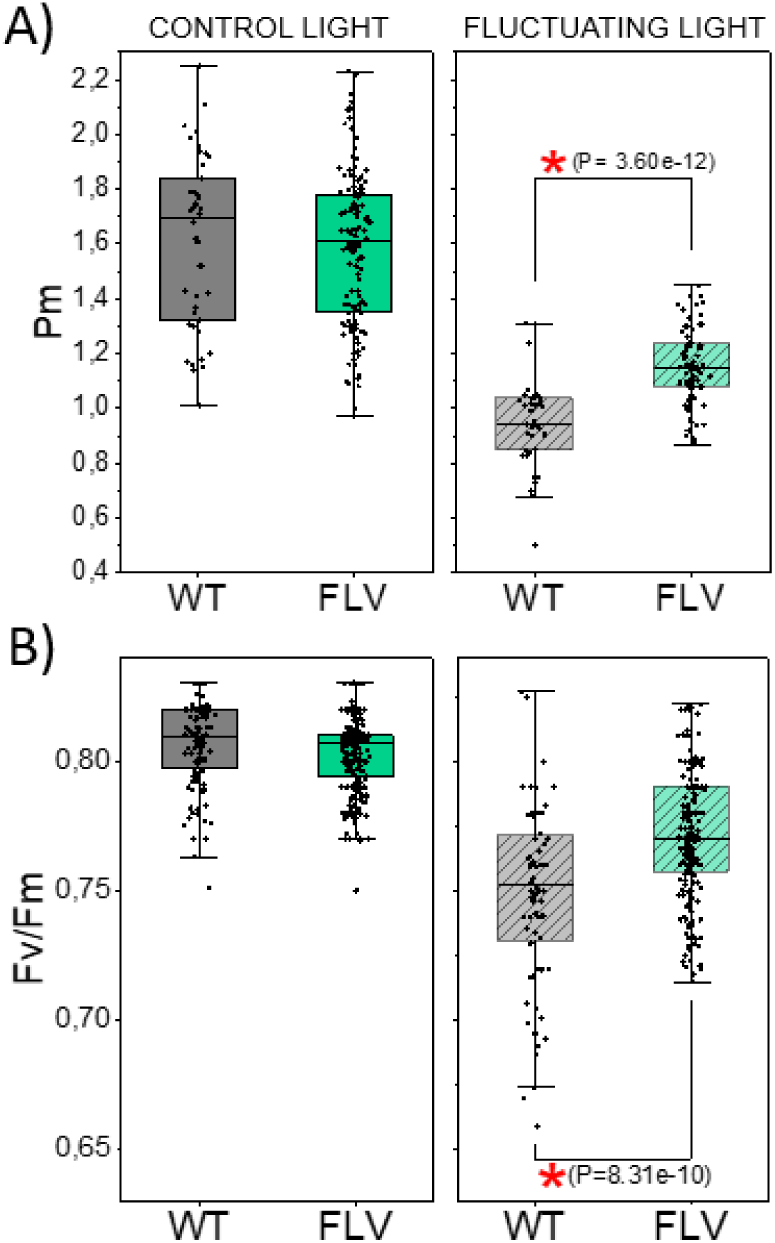
Effect of light regime on PSI and PSII efficiency. Pm (A) and Fv/Fm (B) were measured in dark adapted WT (black) and FLV-expressing lines (green) grown at different light regimes for 14 days. The left panels show data obtained under control light conditions (100 μmol m^-2^ s^-1^; photoperiod 16 hours of light and 8 hours of dark; 25°C). The panels on the right show data for plants grown under fluctuating light conditions (4.5 minutes at 50 μmol m^-2^ s^-1^ followed by 30 seconds at 1000 μmol m^-2^ s^-1^; photoperiod 16 hours of fluctuating light and 8 hours of dark; 16°C). Data was recorded for each plant once a day. (A) WT, n = 3 ± SD. FLV, n = 10 ± SD (control light); WT, n = 3 ± SD. FLV, n = 6 ± SD (fluctuating light). (B) WT, n = 11 ± SD. FLV, n = 25 ± SD (control light); WT, n = 7 ± SD. FLV, n = 18 ± SD (fluctuating light). Differences between WT and mutant plants were examined by one-way ANOVA; a red asterisk indicates statistical significance.

When plants were exposed to fluctuating light conditions (FL; 4.5 minutes at 50 μmol m^-2^ s^-1^ followed by 30 seconds at 1000 μmol m^-2^ s^-1^), all genotypes showed reduced Pm and Fv/Fm values compared to their CL-grown counterparts (Fig. 7). However, FLV expressing lines retained significantly higher Pm and Fv/Fm values as compared to WT plants (Fig. 7), suggesting a photoprotective effect of FLV during plant growth making plants more tolerant to light fluctuations.

## Discussion

### FLV has intrinsic ability to provide burst of electron transport

Upon an abrupt increase of incident light intensity, the synthesis of reducing equivalents and ATP production respond very fast to changes in illumination. This different kinetics can easily result in over-reduction of the electron transport chain, leading to subsequent photodamage (Allahverdiyeva et al. 2013; Gerotto et al. 2016; Chaux et al. 2017; Shimakawa et al. 2017; Bag et al. 2023). The photoprotective effect of FLVs derives from their ability to mitigate imbalances caused by the fast electron transport increase and the slow regulation of energy-consuming metabolic processes. A substantial amount of data supported the idea that FLVs were essential during the dark-to-light transition and during a sudden increase in light intensity to avoid PSI over-reduction. Due to its capacity of rapidly accepting electrons from PSI, FLVs effectively prevent acceptor side limitation and consequent damage with a protective role. This function has been conserved during the evolution of photosynthetic organisms since it was consistently observed across diverse species, including cyanobacteria (Allahverdiyeva et al. 2013), eukaryotic algae (Jokel et al. 2015; Chaux et al. 2017; Burlacot et al. 2018) and non-vascular plants (Gerotto et al. 2016; Shimakawa et al. 2017; Storti et al. 2019, 2020a, 2020b).

In this study, we introduced FLVs from *P. patens* into *N. tabacum* plants, a species widely used for engineering various aspects of photosynthesis. These efforts have included modifications to non-photochemical quenching (Kromdijk et al. 2016), the Rubisco enzyme (Chen et al. 2023b), CO_2_-concentrating mechanisms in chloroplasts (Chen et al. 2023a) and the regulation of photorespiration (South et al. 2019). *P. patens* FLVA/B were expressed and properly accumulated in *N. tabacum* plants (Fig. 1). Proteins were fully active as confirmed by DUAL-PAM measurements of P700 oxidation kinetics (Fig. 1) and acceptor side limitation level (Fig. 4B), and by JTS-100 analyses of electron transport rate (Fig. 6A).

FLV activity monitored at the onset of light activation confirmed that they are fully functional as being able to keep P700 in a safe oxidized state (Fig. 1 and 2) and decreasing PSI acceptor side limitation in respect to WT *N. tabacum* (Fig. 4B). These outcomes are the result of an increased flow of electrons towards FLVs, already detectable 1 second after the light is turned on (Fig. 6A). It is interesting to highlight that after four seconds of illumination there is an inversion and WT plants transport more electrons than FLV-expressing lines (Fig. 6A). These measurements highlighted the complex nature of the competition between CEF and FLV activity, two pathways that compete with natural and artificial electron sinks (Hubáček et al. 2024).

At the onset of light activation, WT *Physcomitrium patens* exhibited a transient increase in ETR. This increase was attributed to FLV activity, as it was not observed in FLV-depleted *P. patens* plants (Gerotto et al. 2016). Already after 6 seconds WT and FLV-depleted *P. patens* showed overlapping electron transport capacity (Gerotto et al. 2016) thus suggesting how FLV presence was negligible. When the same measurements were performed on WT and FLV-expressing *Nicotiana tabacum* plants the results were surprisingly overlapping with *P. patens* ones since FLV-expressing lines showed the transient increase in ETR while steady state ETR was the same in the two lines (Fig. 6). Also, in *Nicotiana tabacum*, FLV contribution appeared minimal at this time point when carbon fixation was fully activated (Fig. 6).

These results confirm that FLV is fully active in angiosperms and that no moss-specific factors are required for its assembly and activity. In particular, the FLV putative electron donors (NADPH and ferredoxin) that have been identified (Peden et al. 2013; Sétif et al. 2020; Beraldo et al. 2024) are clearly present in *N. tabacum* plants. The according results between *P. patens* and *N. tabacum* also suggested that FLV regulation is intrinsic to the protein or simply dependent on some conserved environmental features, such as light intensity or stroma redox potential and does not require any external specific factor.

### FLV competes with CET for electrons

Despite their essential role, FLVs were lost in angiosperms during evolution. It was hypothesized that FLV activity was compensated by an increased CEF and photorespiration activity that ensured enough electron acceptors for PSI (Yamamoto et al. 2016; Hanawa et al. 2017; Fu and Walker 2023). Indeed, FLVs are able to rescue the phenotype of CEF-depleted *A. thaliana* and rice plants without competing with carbon fixation (Yamamoto et al. 2016; Wada et al. 2018; Tula et al. 2020). Accordingly, *P. patens flv* KO lines revealed sustained CEF activity compared to WT plants (Gerotto et al. 2016).

Our ETRI-ETRII data as well as ETR measured in presence of DCMU (Fig. 5C and 6B) showed that CEF in FLV-expressing lines was significantly lower compared to WT thus implying that when FLVs were present and active, they compete for electrons with CEF. Even though earlier experiments highlighted that FLV were able to compensate for CEF removal and that FLV expression diminished electron flow towards CEF, their role could not be considered fully overlapping as WT *N. tabacum* plants exposed to FL revealed significantly lower Pm and Fv/Fm in respect to FLV-carrying lines (Fig. 7).

PCET competes with CET for electrons and, more broadly, FLVs compete with alternative metabolic pathways such as hydrogen production in (Burlacot et al. 2018; Jokel et al. 2019) and also with Mehler reaction in *Chlamydomonas reinhardtii* (Pfleger et al. 2024) as well as nitrogen reduction in the heterocysts of diazotrophic bacteria (Ermakova et al. 2014). This competition is consistent with the complementary role and evolutionary development of increased CET, which compensates for the absence of FLV-mediated photoprotection. The competition between these pathways likely arises from differences in their kinetic properties, with FLVs providing a rapid sink for excess electrons during transient changes in light conditions.

## ACKNOWLEDGMENTS

The author would like to thank Hiroshi Yamamoto and Toshiharu Shikanai for sharing FLVA-2A-FLVB coding sequence.

## Author contributions

**Nicholas Rizzetto**: Investigation, **Eleonora Traverso**: Investigation, Writing – original draft. **Andrea Sabia**: Investigation. **Filippo Fiorin**: Investigation. **Carlotta Francese**: Investigation: Supervision. **Livio Trainotti**: Conceptualization, Supervision. **Tomas Morosinotto**: Conceptualization, Writing – review & editing. **Alessandro Alboresi**: Conceptualization, Supervision, Writing – original draft, Writing – review & editing.

## Data availability statement

The data that support the findings of this study are available from the corresponding author upon reasonable request.

