## Supplemental Figures and Table for "Flavodiiron protein activity outcompetes cyclic electron transport when expressed in angiosperm *Nicotiana tabacum*"

| Clone | Resistance to kanamycin (%; #tested plants) | Presence of transgene (%; #kanamicin resistant plants tested by PCR) | Presence of FLV activity (%; #kanamicin resistant plants tested by P700 oxidation kinetics) |
|---|---|---|---|
| FLV 1 | 66,3%; 191 | 100%; 44 | 93,2%; 44 |
| FLV 7 | 73,1%; 205 | 100%; 19 | 94,7%; 19 |
| FLV 8 | 88,3%; 211 | 100%; 39 | 100%; 39 |

**Table S1. Analysis of T2 generation of FLV transgenic lines.** FLV transgenic lines #1, #7 and #8 were tested for resistance to kanamycin. Resistent lines were then tested for the presence of FLVA gene in the genome and for FLV activity by measurement of redox kinetics of P700 upon dark-to-light transition.

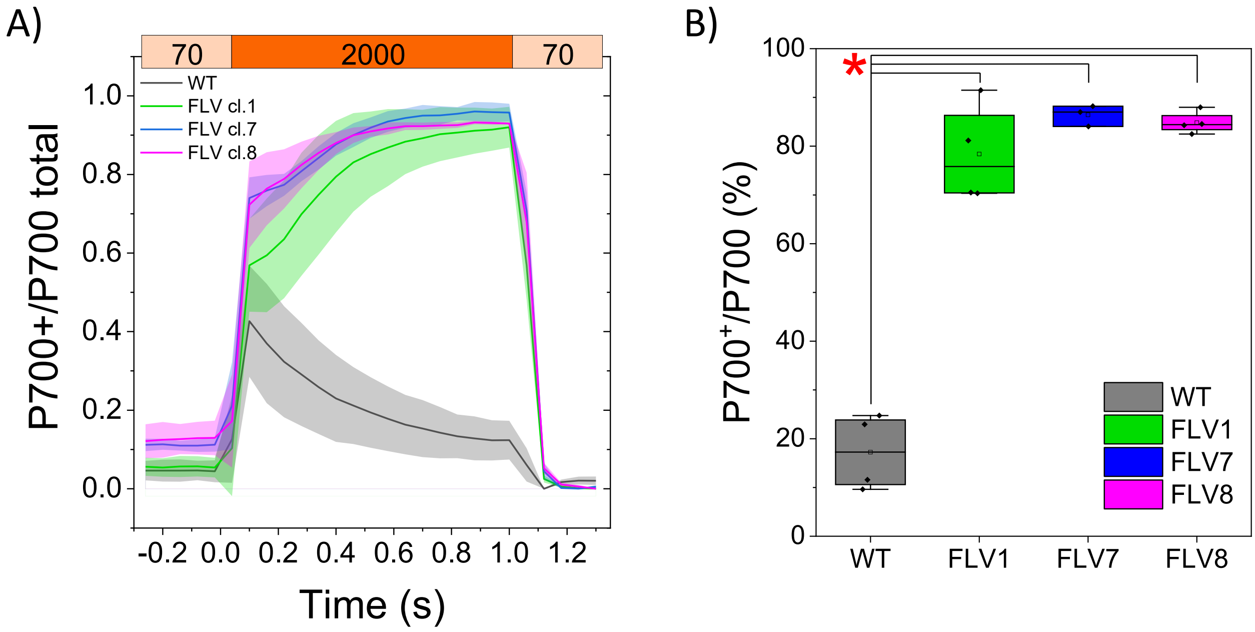


**Fig. S1 Kinetics of oxidized P700 ($P700^+$) in *Nicotiana tabacum* plants.** A) Comparison between wild-type and FLV-expressing lines, the kinetics of oxidized P700 (P700+) during illumination with a short-pulse light (SP: 2000 µmol photons $m^{-2}$ $s^{-1}$, 1 s). Wild-type plants (black) and three FLV-expressing lines (FLV clone 1 in green, FLV clone 7 in blue and FLV clone 8 in magenta) were subjected to SP in the presence of a background light of 70 µmol photons $m^{-2}$ $s^{-1}$. The relative $P700^+$ amount is normalized to Pm, which represents the maximum oxidation level of P700. WT, n = 4 ± SD. FLV cl. 1 n = 4. FLV cl. 7 n = 3. FLV cl. 8 n = 4. A red asterisk indicates statistical significance between WT and all the three FLV-expressing lines, analyzed with one-way ANOVA ($P < 0.01$)

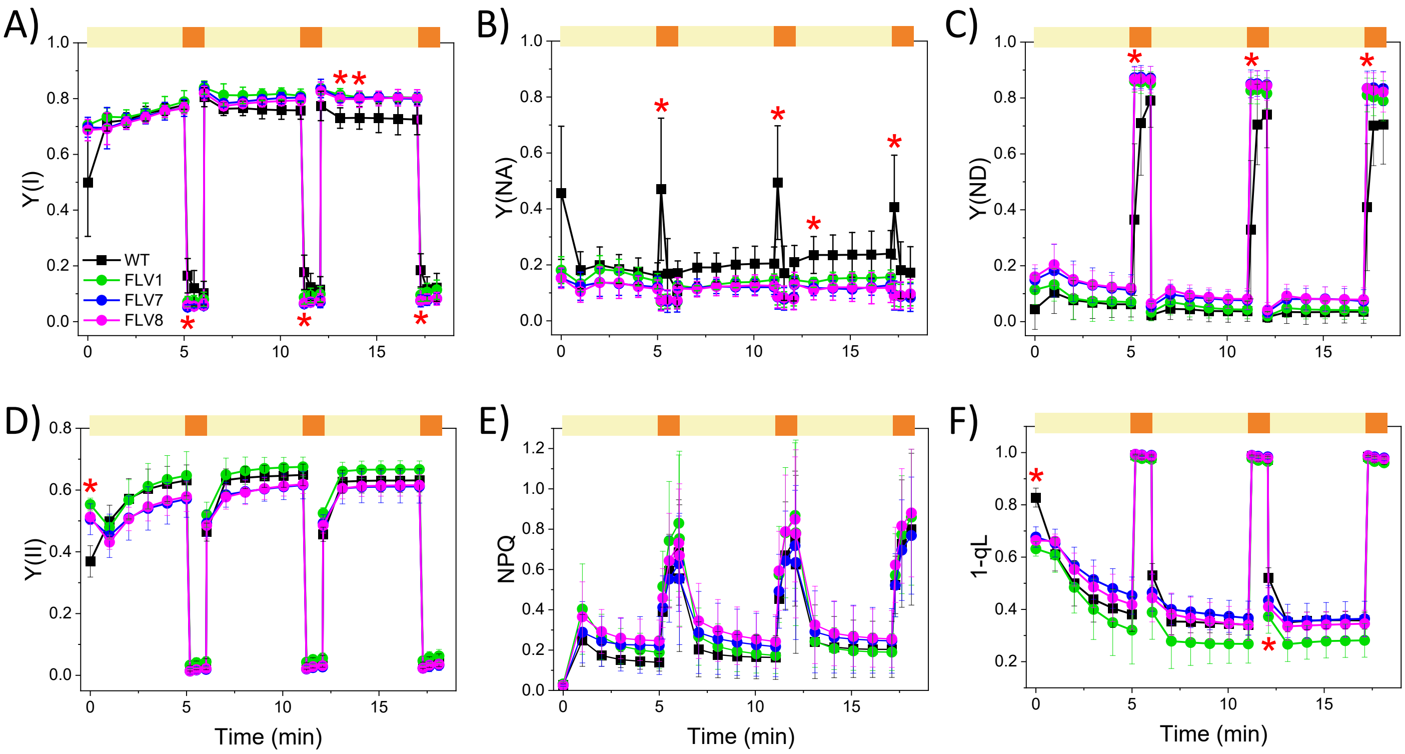


**Fig. S2 *P. patens* FLVs shape electron transport under fluctuating light in *N. tabacum* plants.** Effect of fluctuating light on PSI and PSII: Y(I) (A), Y(II) (B), Y(ND) (C), NPQ (D), Y(NA) (E), and 1–qL (F) in the WT (black squares) and three independent lines expressing FLV proteins (green circles for clone 1, blue circles for clone 7 and magenta circles for line 8). At time 0, after 40 min of dark adaptation, plants were treated with low actinic light (60 µmol photons $m^{-2}$ $s^{-1}$; yellow bars) for 5 min followed by saturating actinic light (1600 µmol photons $m^{-2}$ $s^{-1}$; orange bars) for 1 min. This cycle was repeated 2 more times. Data represent average values ±SD, n=7 for the WT, n=6 for FLV lines clone 1, n=5 for FLV lines clone 7 and n=6 for FLV lines clone 8. Differences between WT and mutant plants in the saturating/limiting light cycles were examined by one-way ANOVA; a red asterisk indicates statistical significance between WT and all the three FLV-expressing lines (P < 0.01).

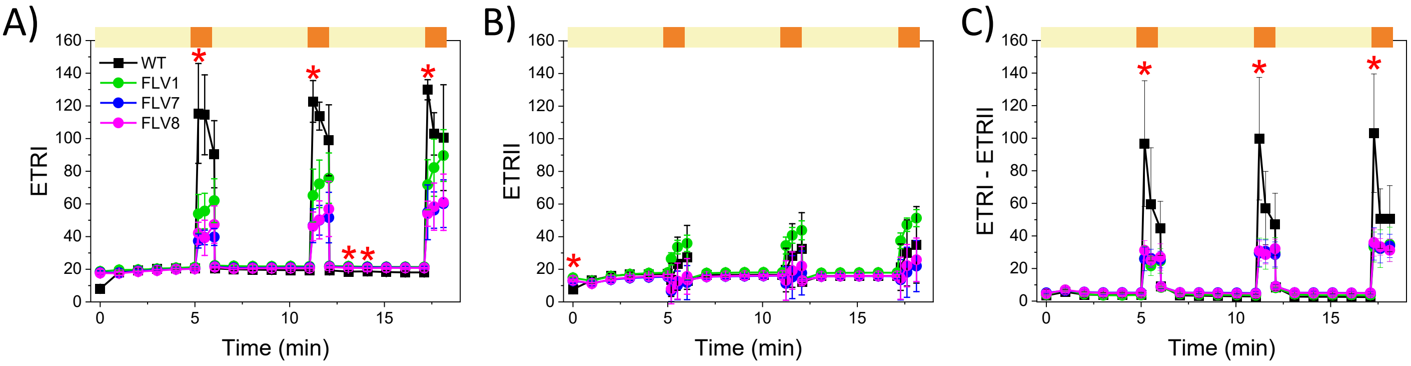


**Fig. S3** ***P. patens*** **FLV repress cyclic electron transport in** ***N. tabacum*** **plants.** Effect of fluctuating light on ETRI (A), ETRII (B) and ETRI-ETRII (C) as a proxy for cyclic electron transport of WT plants (black squares) and two independent lines expressing FLV proteins (green circles for clone 1, blue circles for clone 7 and magenta circles for line 8). At time 0, after 40 min of dark adaptation, plants were treated with low actinic light (60 µmol photons $m^{-2}$ $s^{-1}$; yellow bars) for 5 min followed by saturating actinic light (1600 µmol photons $m^{-2}$ $s^{-1}$; orange bars) for 1 min. This cycle was repeated 2 more times. Data represent average values ±SD, n=7 for the WT, n=6 for FLV lines clone 1, n=5 for FLV lines clone 7 and n=6 for FLV lines clone 8. Differences between WT and mutant plants in the saturating/limiting light cycles were examined by one-way ANOVA; a red asterisk indicates statistical significance between WT and all the three FLV-expressing lines ($P < 0.01$).

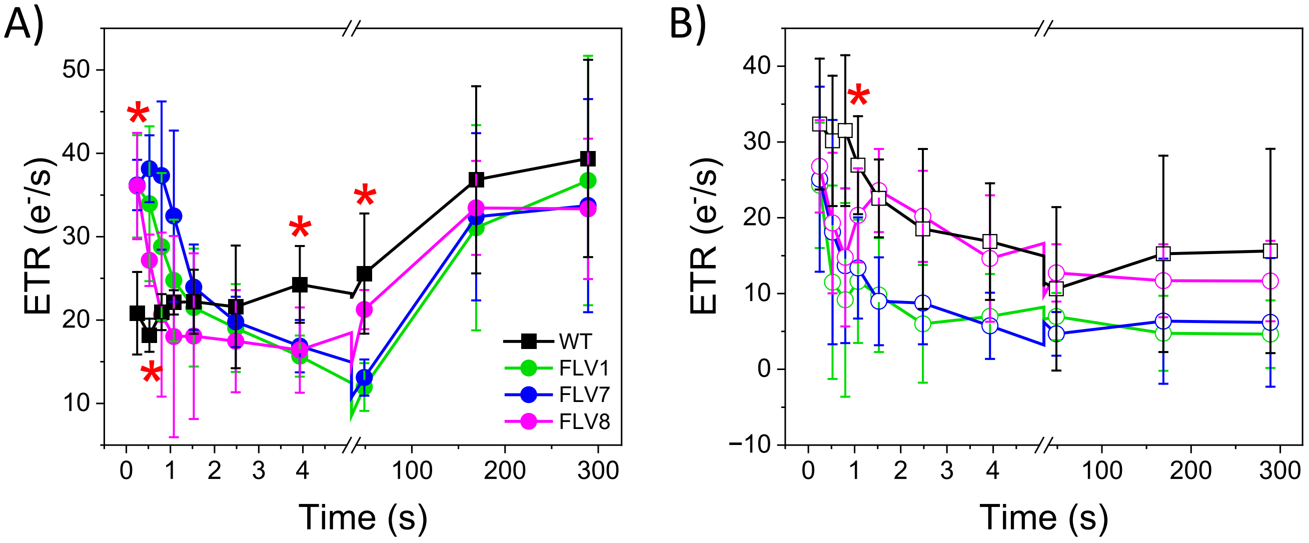


**Fig. S4 Photosynthetic electron transport in *N. tabacum* WT and FLV-expressing lines.** (A) Total photosynthetic ETR measured in WT (black squares) and FLV-expressing lines (green circles for clone 1, blue circles for clone 7 and magenta circles for line 8) at 940 µmol photons $m^{-2}$ $s^{-1}$ actinic light, calculated from electrochromic shift signal. (B) Cyclic electron transport rate measured in the same samples treated with the PSII inhibitor 3-(3,4-dichlorophenyl)-1,1-dimethyl urea (DCMU). Data represent average values ±SD, n=6 for the WT, n=5 for FLV lines clone 1, n=6 for FLV lines clone 7 and n=6 for FLV lines clone 8. Differences between WT and mutant plants were examined by one-way ANOVA; a red asterisk indicates statistical significance ($P < 0.01$).

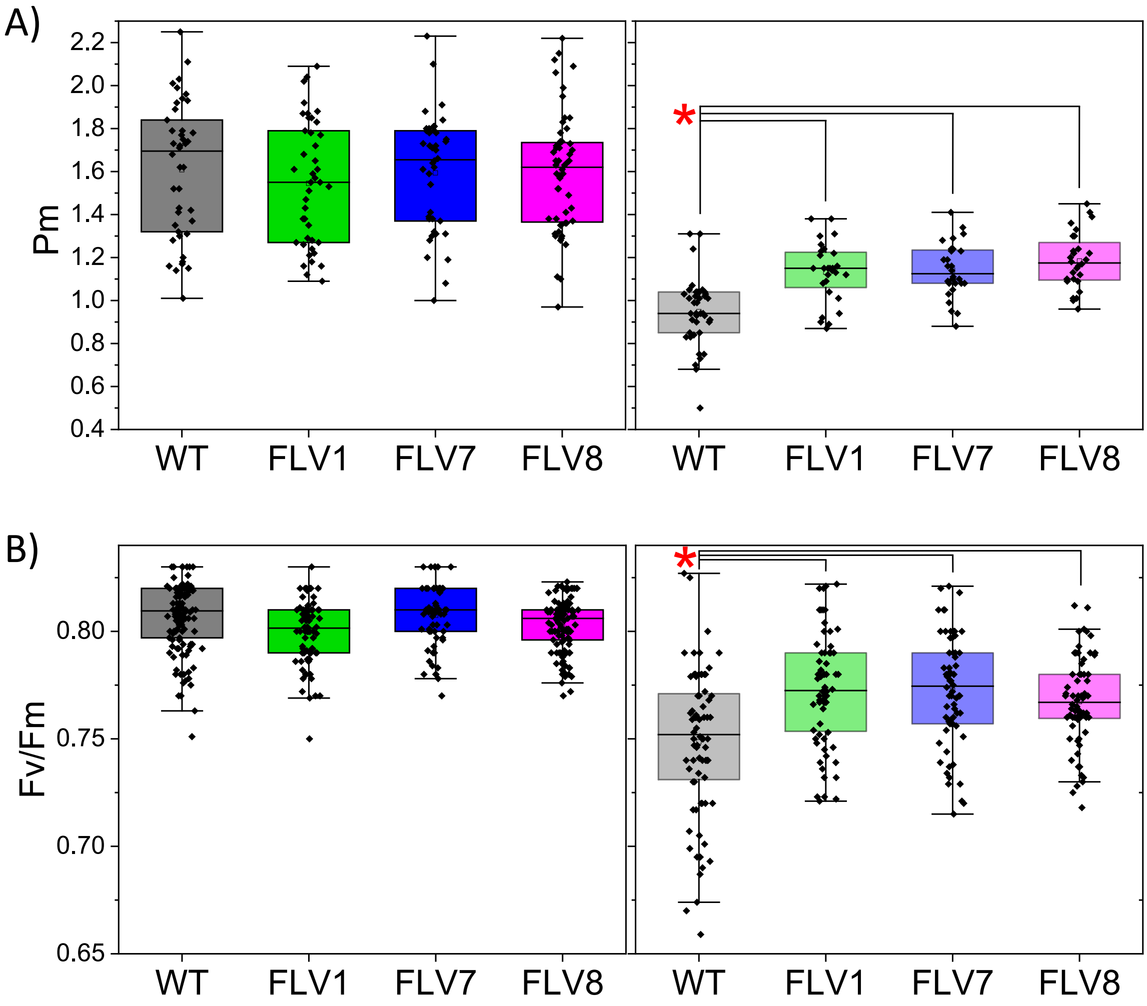


**Fig. S5 Fig. 7 Effect of light regime on PSI and PSII efficiency.** Pm (A) and Fv/Fm (B) were measured in dark adapted WT (black) and FLV-expressing lines (FLV clone 1 in green, FLV clone 7 in blue and FLV clone 8 in magenta) grown at different light regimes for 14 days. The left panels show data for plants grown under standard light conditions (100 μmol photons $m^{-2}$ $s^{-1}$; photoperiod 16 hours of light and 8 hours of dark; 25°C). The panels on the right show data for plants grown under fluctuating light conditions (4.5 minutes at 50 μmol photons $m^{-2}$ $s^{-1}$ followed by 30 seconds at 1000 μmol photons $m^{-2}$ $s^{-1}$; photoperiod 16 hours of fluctuating light and 8 hours of dark; 16°C). Data were recorded for each plant once a day. (A) WT, n=3. FLV, n=3 for clone 1, n=3 for clone 7, n=4 for clone 8 (standard light); WT, n=3. FLV, n=2 for clone 1, n=2 for clone 7, n=2 for clone 8 (fluctuating light). (B) WT, n=11. FLV, n=8 for clone 1, n=6 for clone 7, n=11 for clone 8 (standard light); WT, n=7. FLV, n=6 for clone 1, n=6 for clone 7, n=6 for clone 8 (fluctuating light). Differences between WT and mutant plants were examined by one-way ANOVA; a red asterisk indicates statistical significance ($P < 0.01$)
